# GBAviewer: a structural database of GBA1 variants in Parkinson’s disease

**DOI:** 10.64898/2026.09.05.749596

**Authors:** Andrew N. Bayne, Sitki Cem Parlar, Morvarid Ghamgosar Shahkhali, Lorna Chebon-Bore, Priyabrata Halder, Ziv Gan-Or, Jean-Francois Trempe

## Abstract

**Background:** *GBA1* variants are common risk factors for Parkinson’s disease (PD), yet the structural consequences of most variants remain uncharacterized. No centralized resource currently integrates the growing number of GCase structures, compounds, and PD-associated variants.

**Objectives:** To develop an interactive tool mapping *GBA1*-PD missense variants onto GCase structures, alongside interactors and therapeutic compounds.

**Methods:** We built GBAviewer, an R/Shiny web server integrating GBA1-PD Browser variants with crystal structures, cryo-EM complexes, AlphaFold3 models, and AlphaMissense scores. We also modelled a putative GCase-Saposin C-glucosylceramide ternary complex using AlphaFold3.

**Results:** GBAviewer maps variants in 3D across the GCase active site, LIMP-2 and SapC surfaces, chaperone pockets, and dimerization interface. Our ternary model corroborates SapC binding to the GCase active site entrance, where several variants of unknown significance cluster.

**Conclusions:** GBAviewer is a freely accessible platform for structural analysis of *GBA1* variants, supporting mechanistic studies and therapeutic development. Available at: https://g-can.shinyapps.io/GBAviewer/.

## Introduction

Variants in *GBA1*, encoding the lysosomal enzyme glucocerebrosidase (GCase), are common genetic risk factors for Parkinson’s disease (PD) (1). Biallelic loss of function mutations in *GBA1* lead to Gaucher disease (GD), an autosomal recessive lysosomal storage disorder classically divided into type I (non-neuronopathic), type II (acute neuronopathic), or type III (chronic neuronopathic). *GBA1* variants are typically classified as mild (causing GD type I when biallelic) or severe (causing GD 2/3 when biallelic). A third class consists of risk variants (e.g. pE326K, pT369M) that are associated with increased PD risk without causing GD. Notably, GD patients and heterozygous *GBA1* carriers who do not develop GD still carry an increased risk for PD. In heterozygous carriers, the odds ratios for PD range from ∼2-3 for mild mutations (e.g. p.N370S) to over 5-10 for severe mutations (e.g. p.L444P) (2).

*GBA1*-associated PD (GBA1-PD) is characterized by an earlier age of onset (2-3 years for mild variants or roughly 5 years for severe variants), more rapid motor and cognitive decline, and prominent non-motor symptoms compared with idiopathic PD (3–5). However, the penetrance of *GBA1* variants is low and age-dependent, and the mechanisms linking specific *GBA1* mutations to neurodegeneration remain elusive (6,7).

GCase is a lysosomal hydrolase that degrades glucosylceramide (GlcCer) and glucosylsphingosine (GlcSph), following (8). Saposin C (SapC) facilitates GlcCer hydrolysis, after GCase is trafficked by LIMP-2 from the endoplasmic reticulum (ER) to the lysosome (9,10). *GBA1* variants can impair GCase through reduced catalytic activity or defects in folding, stability, and trafficking, with downstream effects including lipid dysregulation, ER stress, and α-synuclein pathology (reviewed in 11,12). How these mechanisms contribute to GBA1-PD remains under investigation. Still, the GCase pathway has emerged as a therapeutic target in GBA1-PD, with clinical trials spanning pharmacological chaperones (*e*.*g*. ambroxol), allosteric activators, substrate reduction therapy, and gene therapy (12,13)

The GBA1-PD Browser (https://pdgenetics.shinyapps.io/gba1browser/) systematically compiled over 350 *GBA1* variants reported in PD, classifying them as severe, mild, risk variant, or unknown based on available clinical data (14). However, the majority (∼70%) of these variants remain of unknown clinical significance due to their rarity and lack of functional characterization.

To bridge this gap, we developed GBAviewer, an interactive R/Shiny web server built as part of the GBA1-Canada (G-Can) initiative. GBAviewer maps GBA1-PD variants onto high-resolution 3D structures alongside binding partners and small molecules, enabling researchers to rapidly assess the structural context of variants for downstream characterization.

## Methods

### Data Source and Variant Processing

We extracted 351 unique missense single-nucleotide variants (SNVs) from the GBA1-PD Browser (14). Variants retain their American College of Medical Genetics (ACMG) Gaucher disease and GBA1-PD Browser classifications. Variants were processed and annotated with AlphaMissense (AM) pathogenicity scores (15) and DynaMut2-predicted changes in apo-GCase stability (ΔΔG) (16). Variants are drawn from a static snapshot of the GBA1-PD Browser at each release and are manually synchronized. The snapshot date is displayed in the application.

### Structural Aggregation and Modelling

Experimental GCase structures were aggregated from the Protein Data Bank (PDB), including apo GCase, the GCase dimer (8P41), the GCase–LIMP-2 cryo-EM complex (9FJF), and complexes with orthosteric and allosteric small molecules (Table 1). Putative GCase-ambroxol and GlcCer-GCase models were generated using DynamicBind (17). GCase-SapC and GCase-SapC-GlcCer complexes were modelled using a local installation of AlphaFold3 (AF3) (18). Ternary complex modelling used three seeds per prediction (15 models total).

**Table 1.** Summary of GCase structures and binding partners integrated in GBAviewer.

| Structure | Ligand / Partner | PDB | Type | Reference |
| --- | --- | --- | --- | --- |
| **Protein complexes** |  |  |  |  |
| GCase (apo) | — | 2NT0 | X-ray | Lieberman et al. 2007 (25) |
| GCase (apo) | — | — | AlphaFold3 | This study |
| GCase + GCase' | GCase (dimer) | 8P41 | X-ray | Schulze et al. 2024 (26) |
| GCase + LIMP-2 | LIMP-2 | 9FJF | Cryo-EM | Dobert et al. 2025 (20) |
| GCase + SapC | Saposin C | — | AlphaFold3 | This study |
| **Orthosteric ligands** |  |  |  |  |
| GCase + IFG | Isofagomine | 2NSX | X-ray | Lieberman et al. 2007 (25) |
| GCase + ABX | Ambroxol | — | DynamicBind | This study |
| GCase + Cpd33 | Roche Cpd33 | 6T13 | X-ray | Benz et al. 2021 (27) |
| GCase + UCB32 | UCB32 | 8P41 | X-ray | Schulze et al. 2024 (26) |
| **Allosteric ligands** |  |  |  |  |
| GCase + UCB31 | UCB31 | 8P3E | X-ray | Schulze et al. 2024 (26) |
| GCase + Cpd28 | Astex Cpd28 | 9FB2 | X-ray | Palmer et al. 2024 (28) |
| GCase + Cpd24 | Astex Cpd24 | 9FAZ | X-ray | Palmer et al. 2024 (28) |

### Browser Construction

GBAviewer was built in R (v4.4.2) using Shiny and the NGLVieweR package for interactive molecular visualization (19). Variants are displayed as colour-coded residues on 3D structures, filterable by AM score, GBA1-PD classification or ACMG tier. The variant table integrates positional, genetic, and pathogenicity data. GBAviewer is freely available at https://g-can.shinyapps.io/GBAviewer/.

## Results

### GBAviewer integrates variant, structural, and predictive annotations

GBAviewer integrates 351 missense GBA1-PD variants across multiple GCase structures and binding partners into a unified, interactive 3D platform **(Figure 1A,B)**. Users can overlay disease variants onto the GCase structure in the context of the catalytic dyad (Glu235/Glu340), the LIMP-2 trafficking interface, the SapC activation surface, allosteric chaperone binding pockets, and the dimerization interface. Colour-coding and surface displays are modifiable by GBA1-PD clinical type, ACMG classification, or AM pathogenicity score. The accompanying variant table provides searchable, sortable access to genetic and computational annotations for each variant and can be filtered to display only variants at a user-selected residue.

**Figure 1.**
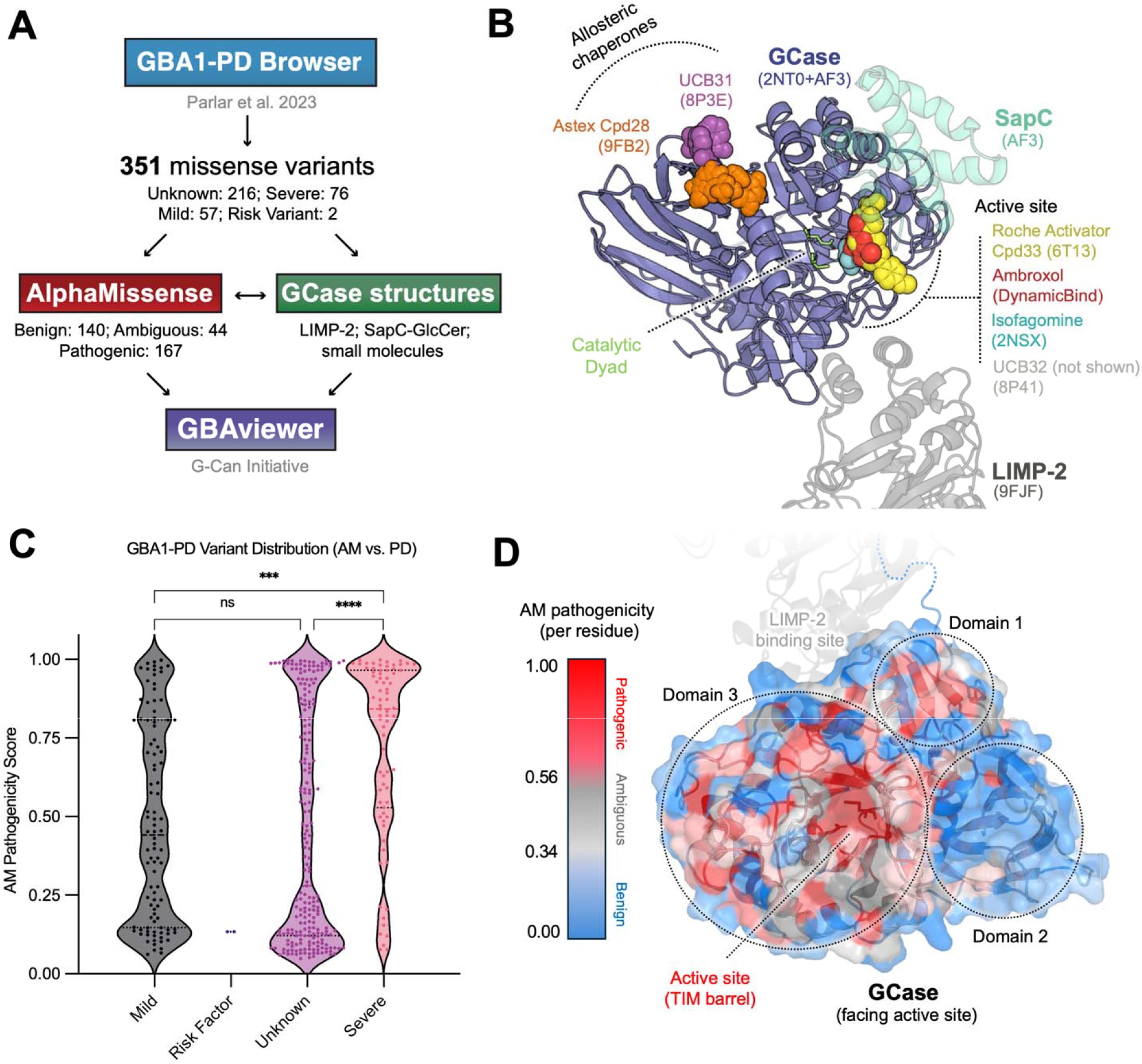
GBAviewer unifies clinical, structural, and predictive data for GBA1-PD variants. **A)** Overview of the GBAviewer pipeline, harmonizing 351 clinical variants from the GBA1-PD Browser with experimental structures, computational models, and AlphaMissense (AM) predictions. **B)** Composite structural model of GCase highlighting key functional interfaces, binding partners, and small molecules integrated within the platform. **C)** Distribution of AM pathogenicity scores across established GBA1-PD clinical classifications from the GBA1-PD Browser classes: mild (GD type I when biallelic), severe (GD type II/III when biallelic), risk variant (raises PD risk without causing GD), and unknown. D) GCase surface coloured by mean per-residue AM pathogenicity, independent of variant presence. Mutation-intolerant regions cluster around the active site and TIM barrel.

To validate the integration of AM scores, we compared AM pathogenicity predictions against both GBA1-PD Browser and ACMG classifications. AM scores were significantly higher for severe versus unknown GBA1-PD variants (p < 0.001) **(Figure 1C)**, suggesting that AM can, on average, discriminate pathogenic variants across our dataset. Mapping the mean per-residue AM pathogenicity onto the GCase structure reveals that mutation-intolerant residues cluster around the active site and TIM barrel **(Figure 1D)**. Still, AM misclassifies ∼20% of disease-causing alleles (severe or mild) as benign, including the well-established p.N370S. This limitation underscores the utility of GBAviewer, in which AM predictions can be directly compared within genetic, biochemical, and structural contexts. Variants are also annotated with published *in vitro* GCase activity measurements.

### A putative structure of the GCase-SapC-GlcCer ternary complex

GCase requires SapC for efficient hydrolysis of GlcCer in the lysosome, yet no experimental structure of the GCase-SapC complex exists. We asked whether co-folding via Alphafold 3 (AF3) could produce a GCase-SapC arrangement consistent with previous biochemical studies (20, 21), and whether a putative ternary GCase-SapC-GlcCer complex could be modelled.

Across AF3 models, SapC occupied a consistent position at the GCase active site entrance **(Figure 2A)**. GlcCer inclusion increased the GCase-SapC interface confidence and SapC pose consistency, with longer GlcCer acyl chains further ordering the interface **(Figure 2B)**. This effect was selective to SapC, as saposins A, B, and D, which activate other lysosomal enzymes, did not show a comparable increase in interface confidence. The modelled interface is also consistent with one of the cross-link sites identified by mass spectrometry involving GCase Lys194 **(Figure 2A)** (20). Notably, the modelled SapC interface overlaps with the GCase dimerization interface, consistent with prior work showing that SapC dissociates GCase dimers (21).

**Figure 2.**
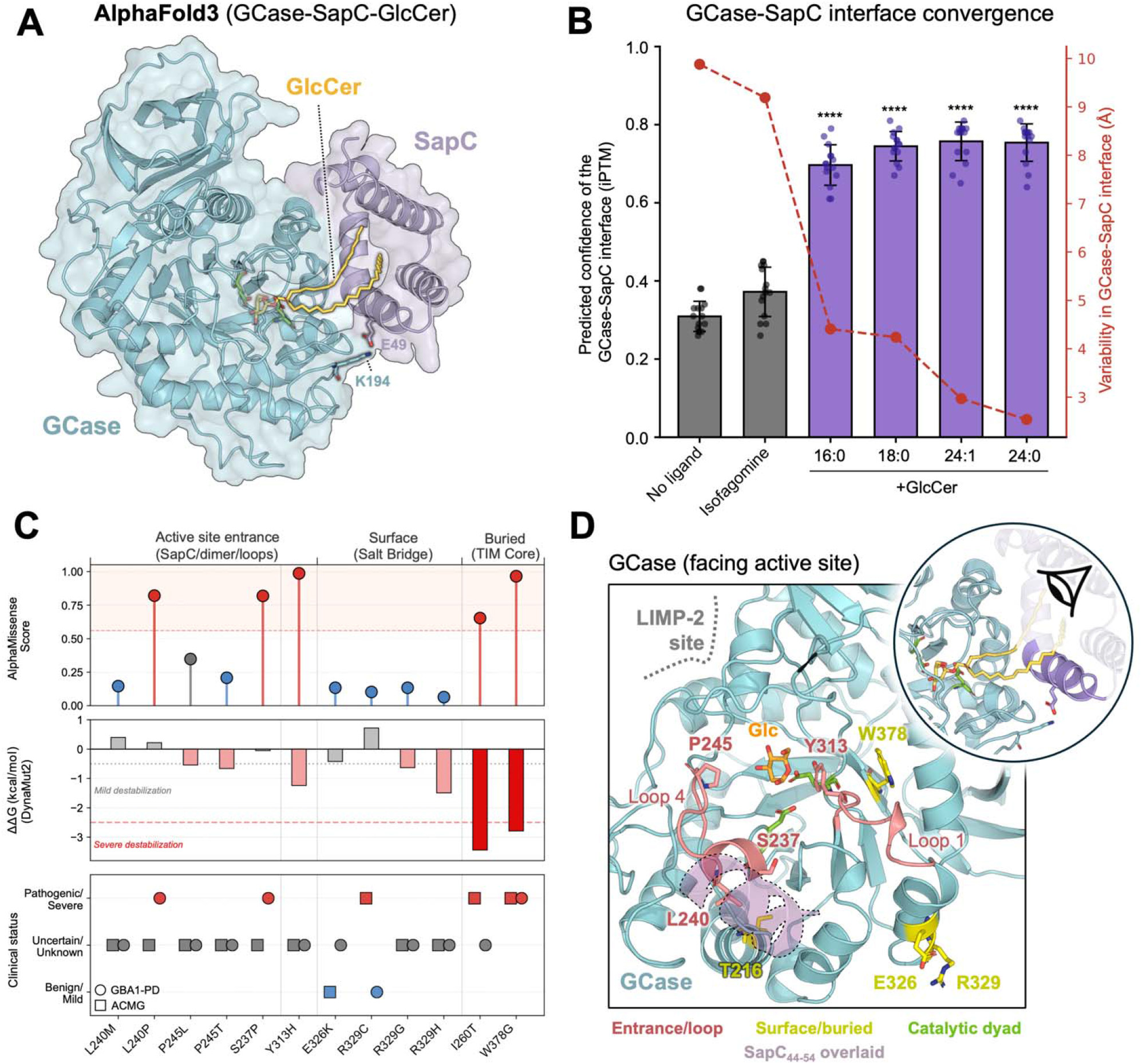
Structural mapping of GCase contextualizes unresolved GBA1-PD variants. **A)** AlphaFold 3 prediction of the putative GCase-SapC-GlcCer (24:0) ternary complex. **B)** Confidence of the GCase-SapC interface across ligand-free (apo), isofagomine-bound, and GlcCer-bound states. Bars show the interface predicted TM score (iPTM) as mean ± SD across 15 models (5 models x 3 seeds). The red dashed line (right axis) indicates structural convergence (mean pairwise Cα RMSD of interface residues after superposition on GCase). Significance was determined via one-way ANOVA with Dunnett’s test against apo (**** p < 0.0001). **C)** AM pathogenicity, DynaMut2 predicted stability change (ΔΔG), and clinical status (ACMG and GBA1-PD classifications) for each highlighted variant. **D)** Highlighted variants mapped onto the ternary model, viewed toward the active-site entrance and coloured by structural context (entrance/loop, surface/buried, catalytic dyad). SapC (purple) is shown as its α3 helix (residues 44-54), the region closest to the active site entrance. GlcCer acyl tails are hidden for clarity. The glucose head group is retained (Glc).

### GBAviewer contextualizes unresolved or rare *GBA1* variants

By delineating functional interfaces, GBAviewer provides structural context for understudied GCase variants. Notably, the active site, SapC binding surface, and dimerization interface, all converge on one face of GCase. Several variants of unknown significance (VUS), such as p.S237P, p.L240P/M, and p.Y313H, cluster within this region **(Figure 2C,D)**. For example, p.S237P and p.L240P/M are located on GCase loop 4, a substrate-gating loop that closely borders the predicted SapC interface. Pro245 locates to the center of this loop close to the active site. While p.P245T is catalytically inactive *in vitro* (22), p.P245L remains uncharacterized. Whether loss of proline alone disrupts loop dynamics or side-chain chemistry also contributes remains to be tested. Deep within the active site, p.Y313H locates to loop 1, forming hydrogen bonds with the catalytic E340 **(Figure 2D)**. Loss of these hydrogen bonds is likely to disrupt catalysis, as corroborated by the highly pathogenic AM score (0.99) **(Figure 2C)**.

GBAviewer also enables the rapid profiling of surface-exposed variants. The common PD risk variant GCase p.E326K has been shown to impair LIMP-2 trafficking of GCase via disruption of a salt bridge with R329, which promotes constitutive dimerization of GCase (23). Our dataset identifies three R329 variants (p.R329C, p.R329G, and p.R329H). Although E326 and R329 do not overlap with the LIMP-2 binding surface, GBAviewer reveals they lie closer to the putative SapC and GCase dimerization interfaces **(Figure 2D)**.

In contrast to the multi-functional dynamics occurring at the GCase surface, GBAviewer readily corroborates disease variants acting via severe folding defects or thermodynamic destabilization. Buried core mutations, including the French-Canadian founder allele p.W378G and deep TIM-barrel substitutions, such as p.I260T, exhibit high AlphaMissense scores (> 0.90) and severe destabilization by DynaMut2 (ΔΔG < - 2.5 kcal/mol), two metrics that can be readily inspected on the platform **(Figure 2C,D)**.

## Discussion

By harmonizing GBA1-PD Browser data (14) with experimental and modelled GCase structures, GBAviewer bridges genetic epidemiology and structural biology. AlphaMissense scores align well for most clinical classifications, supporting their use as a supplementary metric for the ∼70% of GBA1-PD variants currently classified as unknown. In addition, structural mapping of mutation-intolerant hotspots flags specific regions to prioritize for functional validation. Recent *in silico* work substantiates this approach, clustering GCase variants by predicted structural effect, enzymatic activity, and GD severity (24).

A core strength of GBAviewer is the capacity to contextualize variants within functional interfaces, enabling rapid hypothesis generation. For example, structural mapping illustrates how GCase dimers and SapC likely utilize superimposable surfaces, consistent with biochemical evidence suggesting these two assemblies are mutually exclusive. Mapping variants to known allosteric and orthosteric drug-binding pockets can inform the design of personalized therapeutics or targeted clinical studies. While sequence-based algorithms tend to misclassify pathogenic surface variants as benign, GBAviewer provides aggregated context to triage these VUS for downstream characterization.

Several limitations to this database must be noted. GBAviewer relies on static structures and does not capture the conformational dynamics of GCase during folding or trafficking. Our ternary complex model, while supported by cross-linking mass spectrometry, lacks the lysosomal membrane in which SapC acts to present lipids to GCase. Lastly, while structural context can suggest whether a variant is likely to disrupt folding or a functional interface, it cannot yet predict its downstream cellular consequences. GBAviewer therefore generates testable structural hypotheses rather than functional predictions.

In conclusion, GBAviewer provides a freely accessible, centralized platform that empowers clinical geneticists and researchers to rapidly assess the structural landscape of *GBA1* mutations. This resource lays the groundwork for hypothesis-driven functional studies and may inform the rational design of variant-specific therapeutic strategies for GBA1-associated neurodegeneration.

GBAviewer is available at https://g-can.shinyapps.io/GBAviewer/.

## Authors’ Roles

1. Variant discovery and validation: A. Conception, B. Organization, C. Execution;
2. Structural database: A. Conception, B. Organization, C. Execution;
3. Manuscript preparation: A. Writing of first draft, B. Review and editing;

ANB: 2A,2B,2C,3A,3B

SCP: 1A,1B,1C,3B

MGS: 1A,1B,1C,3B

LC-B: 1A,2A,3B

PH: 1A,2A,3B

ZG-O: 1A,1B,1C, 2A, 2B, 3B

JFT: 2A, 2B, 3B

## References

1. Sidransky E, Nalls MA, Aasly JO, et al. Multicenter analysis of glucocerebrosidase mutations in Parkinson’s disease. N Engl J Med 2009;361(17):1651–1661. 10.1056/NEJMoa0901281

2. Gan-Or Z, Amshalom I, Kilarski LL, et al. Differential effects of severe vs mild GBA mutations on Parkinson disease. Neurology 2015;84(9):880–887. 10.1212/WNL.0000000000001315

3. Brockmann K, Srulijes K, Pflederer S, et al. GBA-associated Parkinson’s disease: reduced survival and more rapid progression in a prospective longitudinal study. Mov Disord 2015;30(3):407–411. 10.1002/mds.26071

4. Brockmann K, Srulijes K, Hauser AK, et al. GBA-associated PD presents with nonmotor characteristics. Neurology 2011;77(3):276–280. 10.1212/WNL.0b013e318225ab77

5. Gan-Or Z, Liong C, Alcalay RN. GBA-Associated Parkinson’s Disease and Other Synucleinopathies. Curr Neurol Neurosci Rep 2018;18(8):44. 10.1007/s11910-018-0860-4

6. Straniero L, Asselta R, Bonvegna S, et al. The SPID-GBA study: Sex distribution, Penetrance, Incidence, and Dementia in GBA-PD. Neurol Genet 2020;6(6):e523. 10.1212/NXG.0000000000000523

7. Balestrino R, Tunesi S, Tesei S, et al. Penetrance of Glucocerebrosidase (GBA) Mutations in Parkinson’s Disease: A Kin Cohort Study. Mov Disord 2020;35(11):2111–2114. 10.1002/mds.28200

8. Grabowski GA, Gatt S, Horowitz M. Acid beta-glucosidase: enzymology and molecular biology of Gaucher disease. Crit Rev Biochem Mol Biol 1990;25(6):385–414. 10.3109/10409239009090616

9. Sun Y, Qi X, Grabowski GA. Saposin C is required for normal resistance of acid beta-glucosidase to proteolytic degradation. J Biol Chem 2003;278(34):31918–31923. 10.1074/jbc.M302752200

10. Reczek D, Schwake M, Schröder J, et al. LIMP-2 is a receptor for lysosomal mannose-6-phosphate-independent targeting of beta-glucocerebrosidase. Cell 2007;131(4):770–783. 10.1016/j.cell.2007.10.018

11. Hertz E, Chen Y, Sidransky E. Gaucher disease provides a unique window into Parkinson disease pathogenesis. Nat Rev Neurol 2024;20(9):526–540. 10.1038/s41582-024-00999-z

12. Menozzi E, Toffoli M, Deleidi M, et al. New evidence on the clinical, genetic, and biochemical bases of GBA1-Parkinson’s disease: prospects for treatment. Lancet Neurol 2026;25(6):602–614. 10.1016/S1474-4422(26)00090-6

13. Menozzi E, Schapira AHV. Prospects for Disease Slowing in Parkinson Disease. Annu Rev Pharmacol Toxicol 2025;65(1):237–258. 10.1146/annurev-pharmtox-022124-033653

14. Parlar SC, Grenn FP, Kim JJ, Baluwendraat C, Gan-Or Z. Classification of GBA1 Variants in Parkinson’s Disease: The GBA1-PD Browser. Mov Disord 2023;38(3):489–495. 10.1002/mds.29314

15. Cheng J, Novati G, Pan J, et al. Accurate proteome-wide missense variant effect prediction with AlphaMissense. Science 2023;381(6664):eadg7492. 10.1126/science.adg7492

16. Rodrigues CHM, Pires DEV, Ascher DB. DynaMut2: Assessing changes in stability and flexibility upon single and multiple point missense mutations. Protein Sci 2021;30(1):60–69. 10.1002/pro.3942

17. Lu W, Zhang J, Huang W, et al. DynamicBind: predicting ligand-specific protein-ligand complex structure with a deep equivariant generative model. Nat Commun 2024;15(1):1071. 10.1038/s41467-024-45461-2

18. Abramson J, Adler J, Dunger J, et al. Accurate structure prediction of biomolecular interactions with AlphaFold 3. Nature 2024;630(8016):493–500. 10.1038/s41586-024-07487-w

19. Rose AS, Hildebrand PW. NGL Viewer: a web application for molecular visualization. Nucleic Acids Res 2015;43(W1):W576–579. 10.1093/nar/gkv402

20. Dobert JP, Schäfer J-H, Dal Maso T, et al. Cryo-TEM structure of β-glucocerebrosidase in complex with its transporter LIMP-2. Nat Commun 2025;16(1):3074. 10.1038/s41467-025-58340-1

21. Gruschus JM, Jiang Z, Yap TL, et al. Dissociation of glucocerebrosidase dimer in solution by its co-factor, saposin C. Biochem Biophys Res Commun 2015;457(4):561–566. 10.1016/j.bbrc.2015.01.024

22. Malini E, Grossi S, Deganuto M, et al. Functional analysis of 11 novel GBA alleles. Eur J Hum Genet 2014;22(4):511–516. 10.1038/ejhg.2013.182

23. Davis OB, Kung JE, Davis SS, et al. A Common PD-Risk GBA1 Variant Disrupts LIMP2 Interaction, Impairs Glucocerebrosidase Function, and Drives Lysosomal and Mitochondrial Dysfunction. bioRxiv 2025. 10.1101/2025.08.28.672891

24. Lanore A, Tesson C, Basset A, et al. Classification of GBA1 variants and their impact on Parkinson’s disease: an in silico score analysis. npj Parkinsons Dis 2025;11(1):226. 10.1038/s41531-025-01060-6

25. Lieberman RL, Wustman BA, Huertas P, et al. Structure of acid beta-glucosidase with pharmacological chaperone provides insight into Gaucher disease. Nat Chem Biol 2007;3(2):101–107. 10.1038/nchembio850

26. Schulze M-SED, Scholz D, Jnoff E, et al. Identification of ß-Glucocerebrosidase Activators for Glucosylceramide hydrolysis. ChemMedChem 2024;19(7):e202300548. 10.1002/cmdc.202300548

27. Benz J, Rufer AC, Huber S, et al. Novel β-Glucocerebrosidase Activators That Bind to a New Pocket at a Dimer Interface and Induce Dimerization. Angew Chem Int Ed Engl 2021;60(10):5436–5442. 10.1002/anie.202013890

28. Palmer N, Agnew C, Benn C, et al. Fragment-Based Discovery of a Series of Allosteric-Binding Site Modulators of β-Glucocerebrosidase. J Med Chem 2024;67(13):11168–11181. 10.1021/acs.jmedchem.4c00702

